# A targeted CRISPR-Cas9 mediated F0 screen identifies genes involved in establishment of the enteric nervous system

**DOI:** 10.1101/2023.12.28.573581

**Authors:** Rodrigo Moreno-Campos, Eileen W. Singleton, Rosa A. Uribe

**Author notes:** Corresponding author: Rosa A. Uribe.

## Abstract

The vertebrate enteric nervous system (ENS) is a crucial network of enteric neurons and glia resident within the entire gastrointestinal tract (GI). Overseeing essential GI functions such as gut motility and water balance, the ENS serves as a pivotal bidirectional link in the gut-brain axis. During early development, the ENS is primarily derived from enteric neural crest cells (ENCCs). Disruptions to ENCC development, as seen in conditions like Hirschsprung disease (HSCR), lead to absence of ENS in the GI, particularly in the colon. In this study, using zebrafish, we devised an *in vivo* F0 CRISPR-based screen employing a robust, rapid pipeline integrating single-cell RNA sequencing, CRISPR reverse genetics, and high-content imaging. Our findings unveil various genes, including those encoding for opioid receptors, as possible regulators of ENS establishment. In addition, we present evidence that suggests opioid receptor involvement in neurochemical coding of the larval ENS. In summary, our work presents a novel, efficient CRISPR screen targeting ENS development, facilitating the discovery of previously unknown genes, and increasing knowledge of nervous system construction.

## Introduction

The Enteric Nervous system (ENS) is an extensive and complex network of enteric neurons (EN) and glial cells that inhabit the length of the gastrointestinal tract (GI). The ENS controls inherent GI functions, such as gut motility and intestinal barrier function (Furness, 2008). In humans, the ENS is estimated to contain over 600 hundred million neurons intrinsically located between the two muscular layers of the GI tract (Fleming et al., 2020). The ENS has been referred to as “the first brain”, as it has been hypothesized as the first evolved nervous system in all extant animal species (Furness and Stebbing, 2018). Accumulating evidence has revealed ENS importance as a bridge in the microbiota-gut-brain (MGB) axis by establishing bidirectional connections between Central Nervous System (CNS) and microbiota (Geng et al., 2022; Sharkey and Mawe, 2023).

For ENS to be functional it requires the adequate development, differentiation, and assembly of its many different ENs and glial cells during development. In vertebrates, the ENS is primarily derived from neural crest cells (NCC) (Douarin and Teillet, 1973; Epstein et al., 1994). NCCs are proliferative, highly migratory, and multipotent stem cells that delaminate and migrate from the length of the embryonic neuraxis based on microenvironmental cues. NCCs migrate extensively throughout the developing embryo and differentiate into a multitude of different cell types, such as craniofacial tissues or pigment cells (Martik and Bronner, 2017) NCCs commit to an enteric lineage once they enter the foregut mesenchyme, at which point they are named enteric neural crest cells (ENCCs). ENCCs express a combination of marker genes that encode transcription factors and receptors, including *Sox10*, *Foxd3*, *Phox2b*, *Ret* and *Gfra1* (Nagy and Goldstein, 2017; Rao and Gershon, 2018). From the foregut, ENCCs continue to migrate caudally into distal hindgut, where they differentiate into ENs, which are classified based on a combination of molecular and cellular means (Nagy and Goldstein, 2017; Martik and Bronner, 2017; Rao and Gershon, 2018).

There have been great efforts in studying how the ENS forms during early embryonic development due to the numerous ENS diseases known to afflict children and adults. These diseases include Hirschsprung disease (HSCR), a severe enteric neuropathy marked by absence of ENS in the distal gut, leading to gut dysmotility and/or Megacolon, and affecting 1 in 5000 newborns (Montalva et al., 2023). In addition to HSCR, defective ENS function can cause Esophageal Achalasia, Chronic Constipation, and Gastroesophageal Reflux Disease, affecting adults and children worldwide (Brosens et al., 2016).

Zebrafish has gained prominence as a pertinent vertebrate model for biomedical and ENS research. Zebrafish generate plentiful externally fertilized eggs, develop transparent embryos, and share 70% genetic sequence similarity with humans, among which 84% correlates with known human-associated diseases (Choi et al., 2021). Even though zebrafish ENS is less complex in architecture compared to mammals, lacking one layer of ENs (submucosal plexus) and displaying scattered ENs in contrast with the clustering of ganglia, zebrafish ENS and GI functions are largely conserved with mammals (Ganz, 2018; Kuil et al., 2021). Notably, various stages of zebrafish ENS development, genes, and signaling pathways have been elucidated, contributing to a deeper understanding of the molecular basis of ENS development (Ganz, 2018; Kuil et al., 2021). In zebrafish, ENCCs migrate into and along the developing gut between 32-72 hours post fertilization (hpf), and by 4 days post fertilization (dpf) an ENS network begins to form around the whole length of the GI tract (Baker et al., 2022; Bandla et al., 2022; Harrison et al., 2014; Kuil et al., 2021; Uyttebroek et al., 2010). Thus, its genetic conservation, and rapid, simple ENS development, make zebrafish an attractive animal model for elucidating ENS development.

Recently, integration of single-cell RNA sequencing (scRNA-seq) into the zebrafish model for exploration of ENS development has yielded invaluable insights into what genes are expressed during ENS developmental phases (Howard et al., 2021; Kuil et al., 2023). Concurrently, the successful adoption of CRISPR technology for gene disruption in zebrafish, known for its high efficiency in analyzing phenotypes directly in the injected generation (F0), also known as “crispants”, facilitates swift assessment of candidate genes (Gui et al., 2017; Hwang et al., 2013). This synergy between scRNA-seq and CRISPR technology not only holds promise to enhance our understanding of ENS development, but also allows for the rapid and targeted interrogation of novel candidate genes.

In this study, we employed a targeted F0 CRISPR screen, informed by scRNA-seq data (Howard et al., 2021) of early zebrafish ENS development. We pinpointed twelve genes, that when disrupted, led to various ENS development phenotypes across different F0s, or crispants. In particular, we discovered that crispant fish targeting genes encoding for opioid receptors, *oprl1* and *oprd1b,* presented with severe ENS development defects, in which further phenotyping showed reduced ENCC numbers resident along the gut. Subsequently, *oprl1* and *oprd1b* crispant larvae displayed alterations in EN neurochemical coding during ENS maturation. Our subsequent focused investigations of the opioid pathway affirmed its pivotal role in ENS establishment along the gut length, whereby temporal opioid pathway inhibition reduced ENCC abundance along the gut.

## Results

### Construction of an *in vivo* ENS-focused F0 CRISPR screen

Leveraging scRNA-seq datasets is an attractive method for uncovering novel genes expressed during ENS development. Candidate genes can then be targeted with CRISPR gene editing to aid us in understanding their potential functional roles during ENS development. To that end, we focused on our prior zebrafish embryo-to-larval stage single-cell atlas that contained *sox10*:GFP-expressing and -derived cells (Howard et al., 2021) for further analysis. Previously, cell clusters from the 68-70 hpf *sox10*:GFP dataset that captured neuronal populations based on the combinatorial expression of enteric neuron markers such as *elavl3, phox2bb, ret* and *gfra1a* (**Supplementary figure 1A**), were subset and re-clustered, yielding five sub-cluster populations (0-4) (**Figure 1A**) (Howard et al., 2021). Functional enrichment and interactome analysis (Zhou et al., 2019) identified cellular and signaling pathways related to neurons such as membrane trafficking, neuronal system and axon guidance (**Supplementary figure 1B-D**). A total of twelve genes, with high cell-expression distribution from sub-cluster 3 (**Figure 1B**), and which were associated with various predicted neuronal functions, such as receptors, neuropeptides and transcription factors (**Supplementary Figure 1B-D**), were selected for reverse genetic analysis to elucidate their functional significance (**Table 1**). These genes were also present in the protein-protein interaction (PPI) network of sub-cluster 3 in STRING (**Supplementary figure 1E**, **Supplementary Data 1**) (Szklarczyk et al., 2023).

**Figure 1.**
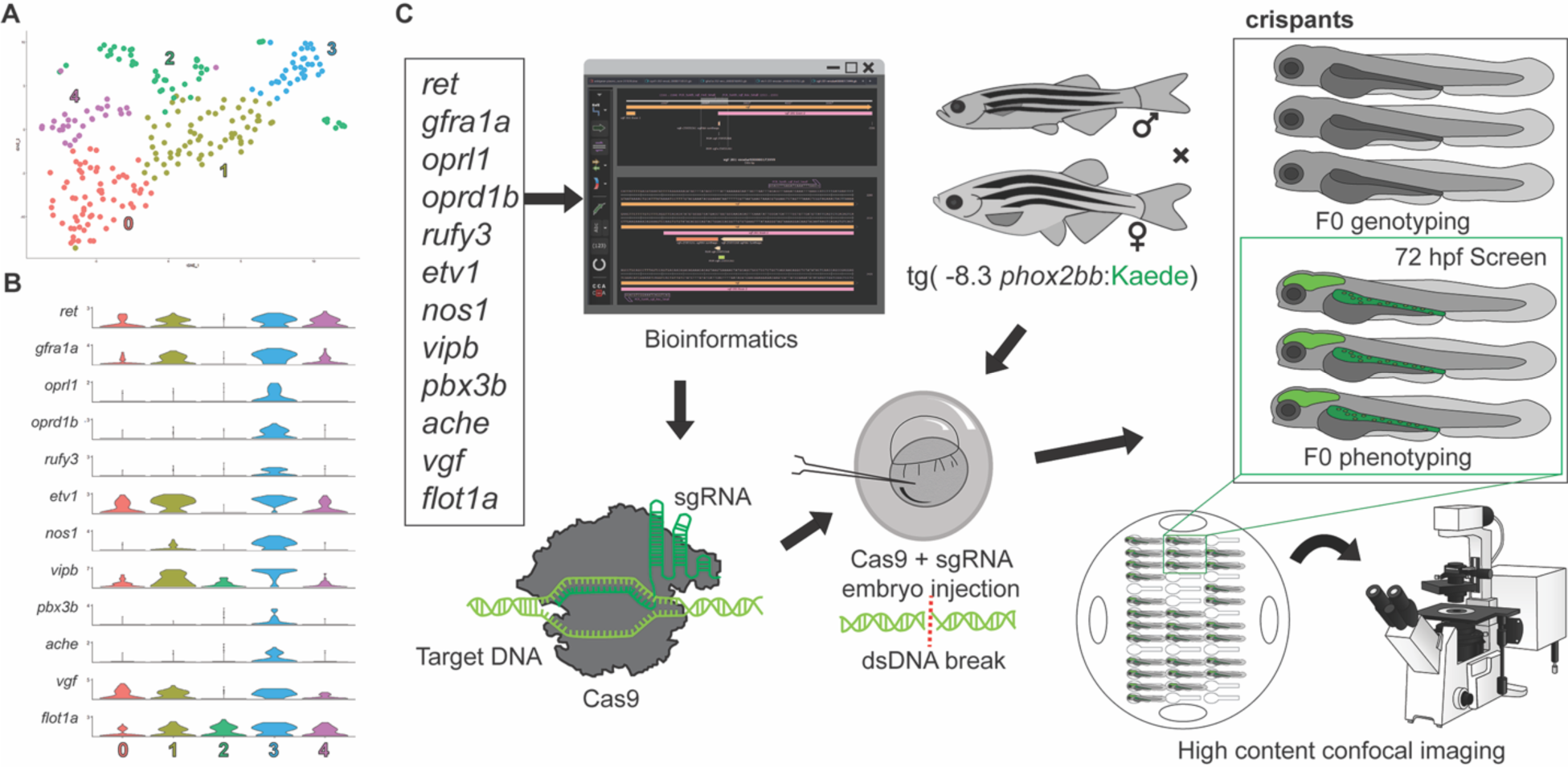
Construction of an F0 CRISPR screen for ENS development. (A) tSNE plot shows five distinct sub-clusters after the subset analysis and re-clustering of Clusters 5 and 12 from the (Howard et al., 2021) 68-70 hpf data set. (B) Violin plots reveal high single-cell expression distribution in sub-cluster 3 of ENS candidate genes. (C) Twelve genes underwent a comprehensive CRISPR screen, involving bioinformatic design, and CRISPR-Cas9 mutagenesis in -8.3*phox2bb*:Kaede zebrafish larvae to visualize enteric cells. The screening strategy included subsequent genotyping validation and high content phenotyping.

**Table 1.**
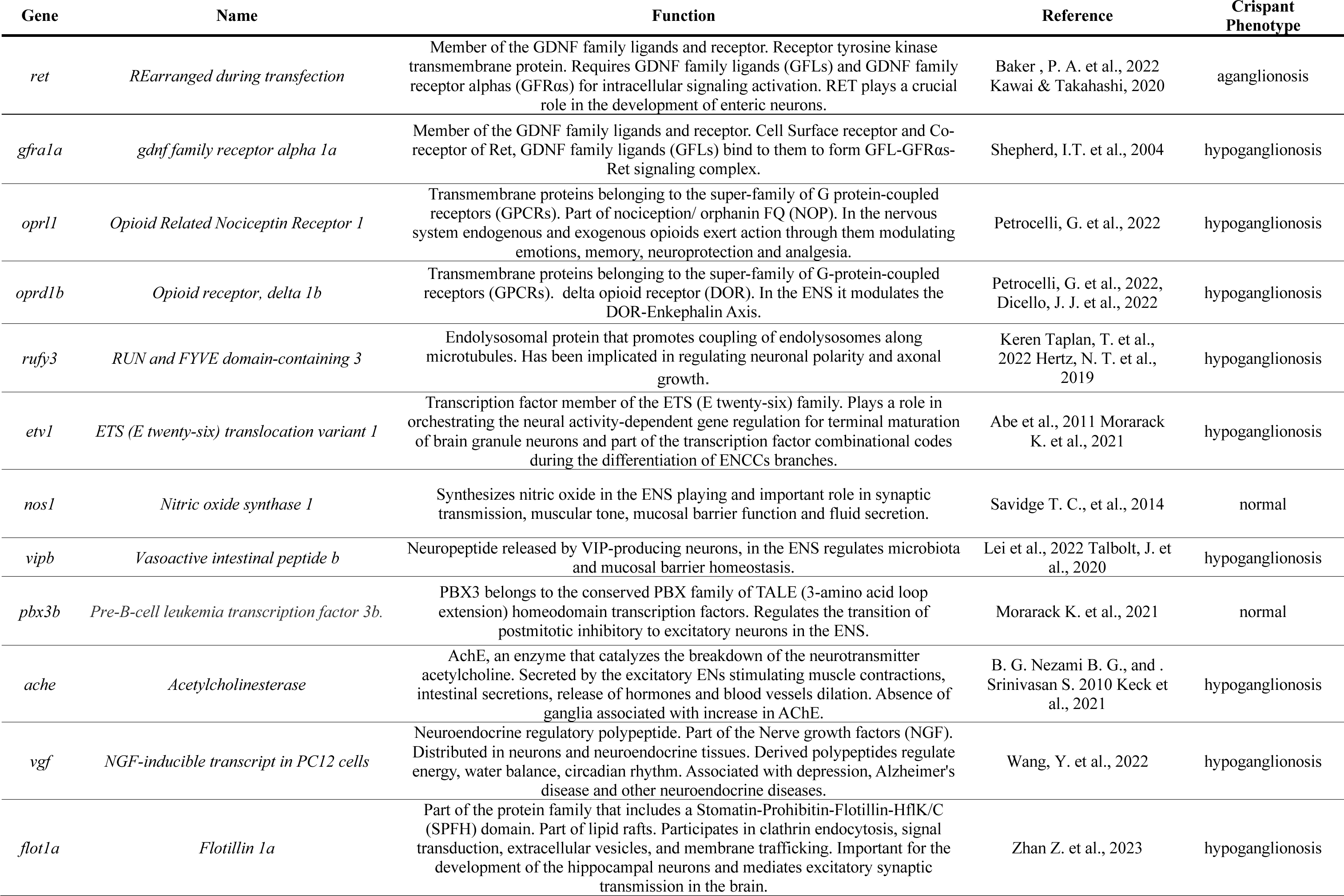
Nomenclature, function and crispant ENS-associated phenotypes from genes of the CRISPR screen.

Next, the candidate ENS genes were input into a targeted ENS F0 CRISPR screen (**Figure 1C**). The screening strategy comprised sequential steps, with each experimental phase requiring confirmation from the preceding one to ensure the integrity and completion of the entire screen. This involved not only genotyping validation and high-content phenotyping, but also a meticulous validation process at each juncture to maintain methodological robustness to uncover phenotypic alterations during ENS development (**Figure 1C**). Leveraging Tg(*-8.3phox2bb*:Kaede) transgenic zebrafish to fluorescently identify ENCCs and ENs (Harrison et al., 2014), we implemented our research strategy by injecting single guide RNA (sgRNA) in complex with Cas9 protein into 1-cell stage transgenic embryos targeting each specific gene (**Supplementary Table 1**), and phenotyping at 72 hpf when the ENS is undergoing neurogenesis. Each of the experimental sets of injected CRISPR F0 embryos, “crispants”, were subjected to genotyping validation and phenotype determination, as outlined in Figure 1C, and described below.

### Candidate gene crispants have ENS genotypic and phenotypic alterations

Genotyping subsets of the crispants for each specific targeted gene via T7 endonuclease 1 (T7E1) mismatch assay was used to detect indel presence, and to enable downstream phenotyping assays (**Figure 2A**) for each batch of F0s. Specifically, our experiments detected indels in a high percentage of embryos (**Figure 2B, C**). This validation assured us that the remaining crispants from each injection pool could be examined for downstream phenotypic ENS alterations.

**Figure 2.**
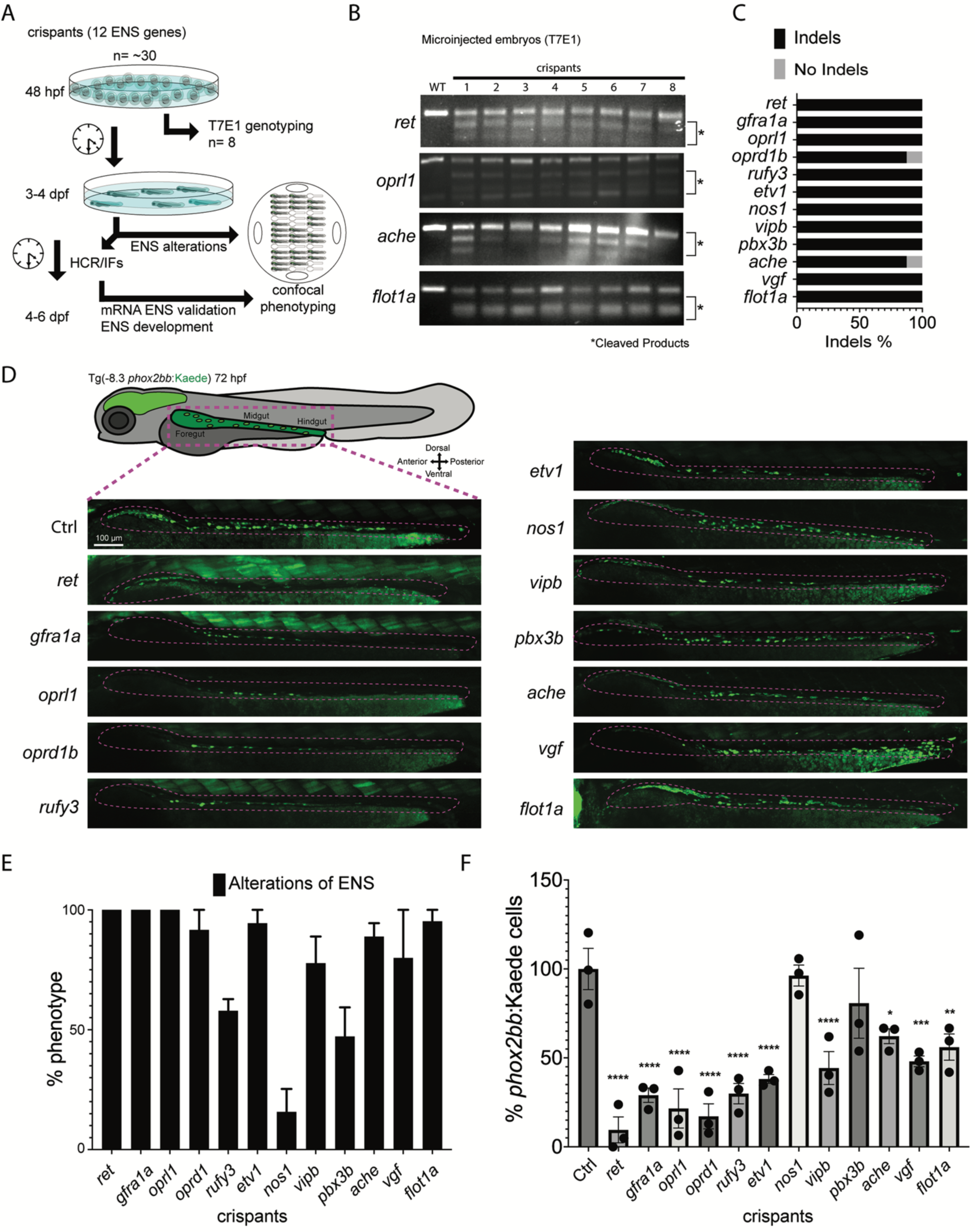
Candidate genes targeted in an F0 CRISPR screen display ENS phenotypic alterations. (A) At 48 hpf, eight crispant embryos from pools of around thirty embryos were used for validation of CRISPR activity via T7E1, for each gene targeted. If the majority had indels then subsets of the pool was grown at 3 to 4 dpf to phenotype their ENS. The phenotyping process was able to combine crispants of different genes by using an agarose cast that enabled the high content semi-automated confocal imaging. An additional fraction of the crispants was analyzed at 4-6 dpf for additional HCR validation or for late phenotypic alterations. (B) Representative images of different T7E1 assays demonstrate indels of different embryos in the specific gene targeted regions. The asterisks denote the presence of cleaved products. (C) Percentage of embryos with CRISPR/Cas9 induced indels of different ENS genes (≥ 2 experiments). (D) Confocal images of Tg(-8.3*phox2bb*:Kaede) ENCCs/ENs for different crispants along the gut at 72 hpf. ENCCs/ENs of the majority of crispants failed to localize distal hindgut. (E) Pools of the twelve ENS genes crispants showing the percentage of phenotypic alterations. (F) Percentage of fluorescent ENCCs and/or ENs along the gut from the different gene crispants. Comparing the mean of the control with the mean of each gene, ANOVA P value ****: <0.0001, ***: 0.0004, **: <0.003, *: <0.05.

To identify ENS phenotypic alterations in the -8.3*phox2bb*:Kaede crispants, we first qualitatively performed our CRISPR screen by examining the colonization success of ENCCs/ENs along the developing gut at 72 hpf. As positive controls for this screen, we utilized sgRNAs against the tyrosine kinase receptor gene, *ret* (REarranged during Transfection), and the GDNF family receptor alpha-1 gene, *gfra1a* (**Figure 2D**). As previously reported, *ret* and *gfra1a* loss-of-function larvae display aganglionosis and hypoganglionosis phenotypes, respectively (Baker et al., 2022; Bandla et al., 2022; Heanue et al., 2016; Shepherd et al., 2004), with aganglionosis presenting as near complete loss of ENs, while hypoganglionosis presents with reductions. Hypoganglionosis alterations were identified in crispants from eight of the screened candidate genes. These genes included: the opioid receptor encoding genes *oprl1* and *oprd1b;* the endolysosomal coupler RUN and FYVE domain-containing protein encoding gene, *rufy3*; ETS (E twenty-six) Variant Transcription Factor 1 encoding gene, *etv1*; vasoactive intestinal peptide encoding gene, *vipb*; Acetylcholinesterase encoding gene, *ache*; VGF Nerve Growth Factor Inducible encoding gene, *vgf*; and the membrane associated protein Flotillin 1 encoding gene, *flot1a.* The genes that didn’t show overt phenotypic alterations in colonization were the nitric oxide synthase encoding gene, *nos1,* and the Pre-B-cell leukemia transcription factor 3b encoding gene, *pbx3b*. The percentage of crispants for each gene with ENS phenotypic alterations was over 80% with all genes, except for *rufy3*, *nos1* and *pbx3b* (**Figure 2E**). Further phenotypic examination of -8.3*phox2bb*:Kaede^+^ larvae crispants for the different candidate genes showed significant reductions in the number of ENs along the gut for all genes tested, when compared with controls, except for *nos1* and *pbx3b* (**Figure 2F**). Overall, our phenotypic results demonstrate that the majority of the CRISPR screened genes are important for ENS establishment, suggesting they may be regulators of ENS formation in zebrafish.

### Genes from the ENS CRISPR screen are expressed along the gut during enteric neurogenesis stages

In order to assay the expression patterns of ENS candidate genes, we performed Hybridization Chain Reaction (HCR) (Ibarra-García-Padilla et al., 2021) with probes specific for select gene transcripts in Tg(-8.3*phox2bb*:Kaede) larvae at 96 hpf, such as *ret*, *etv1*, *oprl1* and *oprd1b* (**Figure 3**). In wholemount, we observed specific expression patterning for *ret*, *oprl1*, *etv1,* and *oprd1b,* as well as *elavl3,* in different regions of the brain and/or the spinal cord (**Supplementary figure 2**). Gene expression in the developing brain has been consistently reported for genes such as *gfra1a* and *elavl3* (Khuansuwan et al., 2019; Shepherd et al., 2004). Previously, we observed expression of *oprl1* within enteric neurons at 70 hpf (Howard et al., 2021). As expected, *ret*, *etv1*, *oprl1* and *oprd1b* were present along the gut ENs (**Figure 3A, C, E, G**), with colocalization seen among the Kaede labeled cells (**Figure 3B-B’’, D-D’’, F-F’’, H-H’’**).

**Figure 3.**
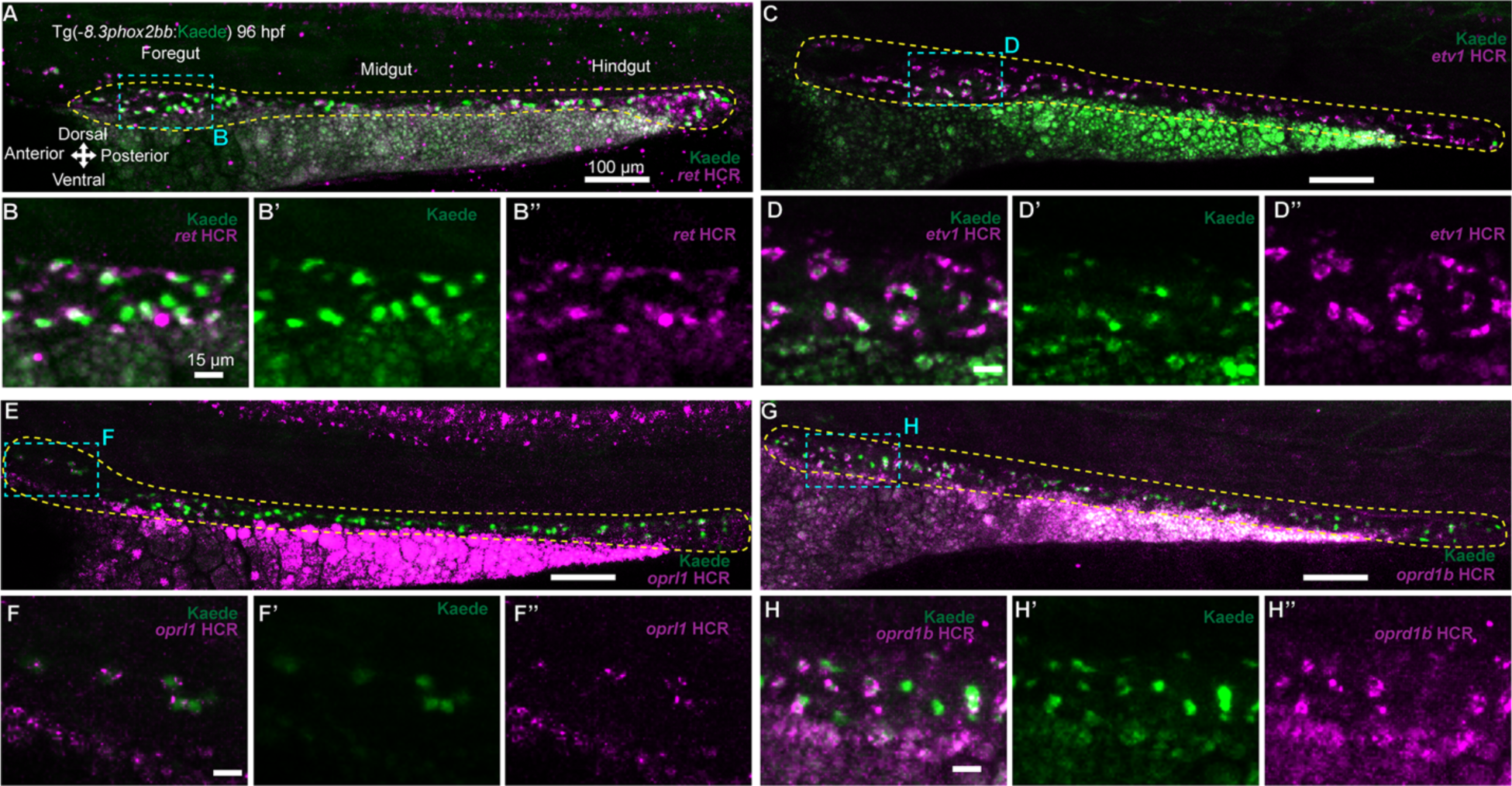
Expression pattern of CRISPR screen selected genes along the ENS during development. (A, C, E, G) Confocal images show HCR-assayed expression for *ret*, *etv1*, *oprl1* and *oprd1b* through the gut of Tg(-8.3*phox2bb*:Kaede) larvae at 96 hpf, dashed yellow lines surround the gut. (B-B”, D-D”, F-F”, H-H”) Magnified regions of the foregut showing colocalization of *ret*, *etv1*, *oprl1* and *oprd1b* (magenta) with ENCCs expressing the Kaede protein (green).

### Chemical inhibition corroborates opioid pathway involvement during ENS development

The opioid receptors have been extensively studied for their physiological roles, and as a pharmacological mechanism for pain treatment in the adult GI tract (Galligan and Sternini, 2017; Wood and Galligan, 2004). Our efforts to locate literature pertaining to opioid receptor involvement in ENS development proved elusive. To further investigate our screen hits, we focused our efforts into chemically targeting the protein products of the opioid receptor encoding genes, the nociception receptor (NOP) Oprl1 (opiate receptor-like 1) or the delta opioid receptor (DOR) Oprd1 (opioid receptor, delta 1a), as crispants for these genes exhibited severe ENS loss in our screen (**Figure 2**), suggesting functional roles for the opioid pathway during ENS establishment. To that end, we employed pharmacological assays using different opioid inhibitors coupled with ENS differentiation assays in -8.3*phox2bb*:Kaede embryos, starting the treatment at 48 hpf for a duration of 48 hours (**Figure 4A**). To inhibit Oprl1 or Oprd1, we treated embryos with different antagonists: LY2940094 and curcumin target Oprl1, and the synthetic peptide agonist DADLE, targets Oprd1b (Mullick et al., 2017; Seo et al., 2018; Statnick et al., 2016): From all treated conditions, when compared with DMSO-treated controls, the incubated larvae displayed hypoganglionosis (**Figure 4B-E**). These data corroborate a role for the opioid receptors during ENS development. When combined with crispant phenotypic data for *oprl1* and *oprd1b* (**Figure 2**), these results suggest that the opioid pathway is required for ENS establishment.

**Figure 4.**
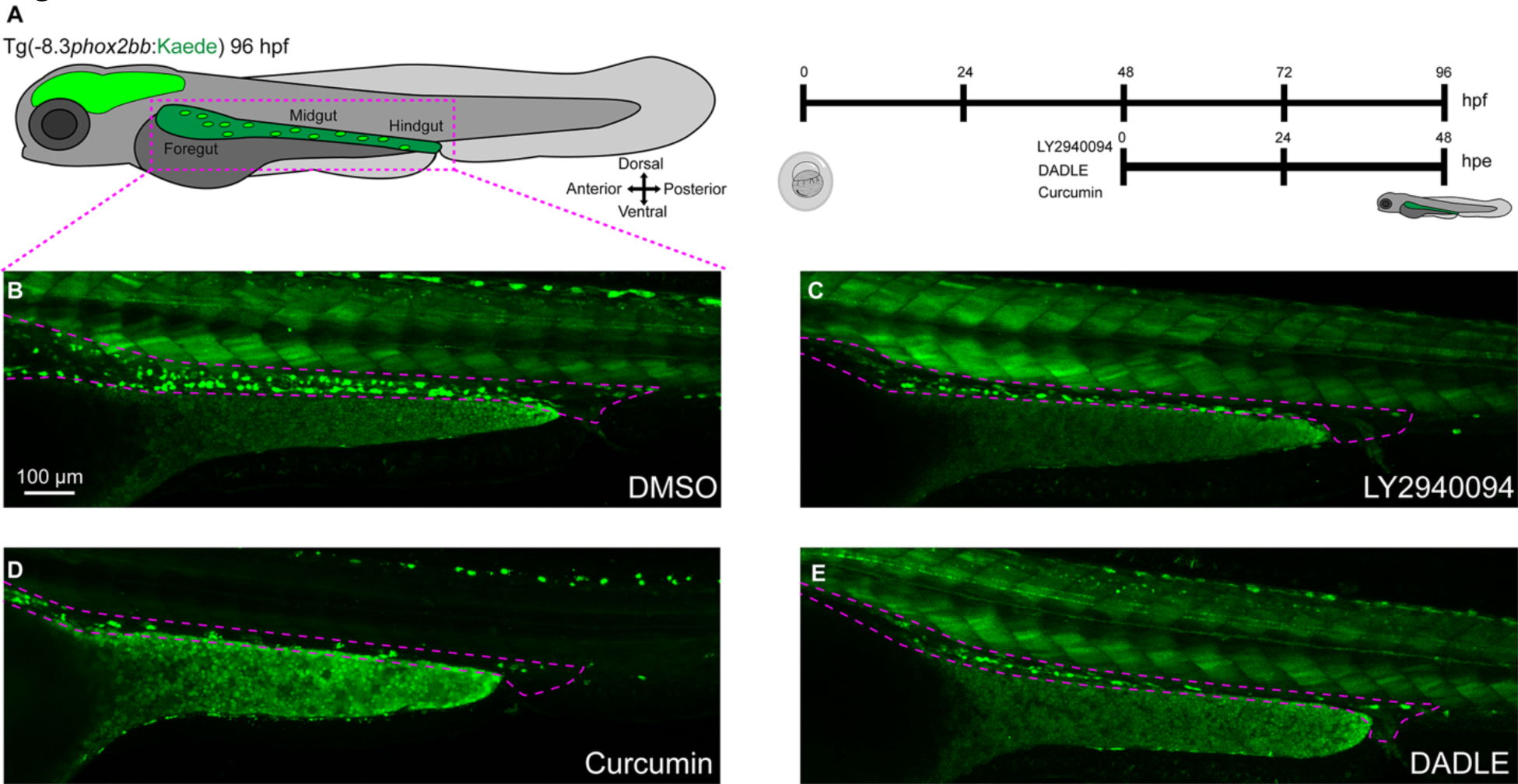
Temporal chemical inhibition of opioid receptors induces ENS developmental defects in zebrafish larvae. (A) Tg(-8.3*phox2bb*:Kaede) embryos were exposed at 48 hpf for 48 hpe (hours post exposure) with the opioid inhibitors, LY2940094, curcumin and DADLE, all of them at 10 µM. (B, C, D, E) Confocal images reveal fluorescent labeled ENCCs/ENs along the gut in larvae that were treated with DMSO, LY2940094, curcumin and DADLE, respectively. Inhibitor-treated larvae show a reduction of Kaede^+^ cells, compared with the DMSO control. Dashed purple lines surround the gut.

### Opioid gene crispants have neurochemical coding alterations in the ENS during development

We next wished to examine if the *oprd1b* and *oprl1* crispants displayed ENS phenotypes later during enteric neuronal differentiation stages. We therefore performed wholemount immunohistochemistry at 6 dpf to detect if changes were present in the neurochemical coding of the ENS. To achieve this, we used antibodies against Phox2b (Howard et al., 2022), and different markers that are present in differentiated ENs, such as HuC/D (Elavl3/4), 5-HT (5-hydroxytryptamine) and Chat (acetylcholine) (Uyttebroek et al., 2010). Control larvae displayed a complete ENS along the gut, and based on the markers we assayed, showed 5 main populations: Phox2b^+^/HuC/D^+^; Phox2b^+^/HuC/D^+^/5-HT^+^; HuC/D^+^/5-HT^+^/Chat^+^; Phox2b^+^; and HuC/D^+^ (**Figure 5A**). Interestingly, while *oprd1b* and *oprl1* crispants generally displayed hypoganglionosis, when compared with control, they both had predominant presence of HuC/D^+^ ENs and a relatively smaller number of HuC/D^+^/5-HT^+^/Chat^+^ and Phox2b^+^/HuC/D^+^ ENs (**Figure 5C, E**). Despite the altered neurochemical code, these ENs were able to populate the hindgut. Overall, these imaging data indicate that, when compared with control, the *oprd1b* and *oprl1* crispants display severe ENS neurochemical coding alterations, suggesting that the opioid pathway regulates the proper establishment of ENS during development.

**Figure 5.**
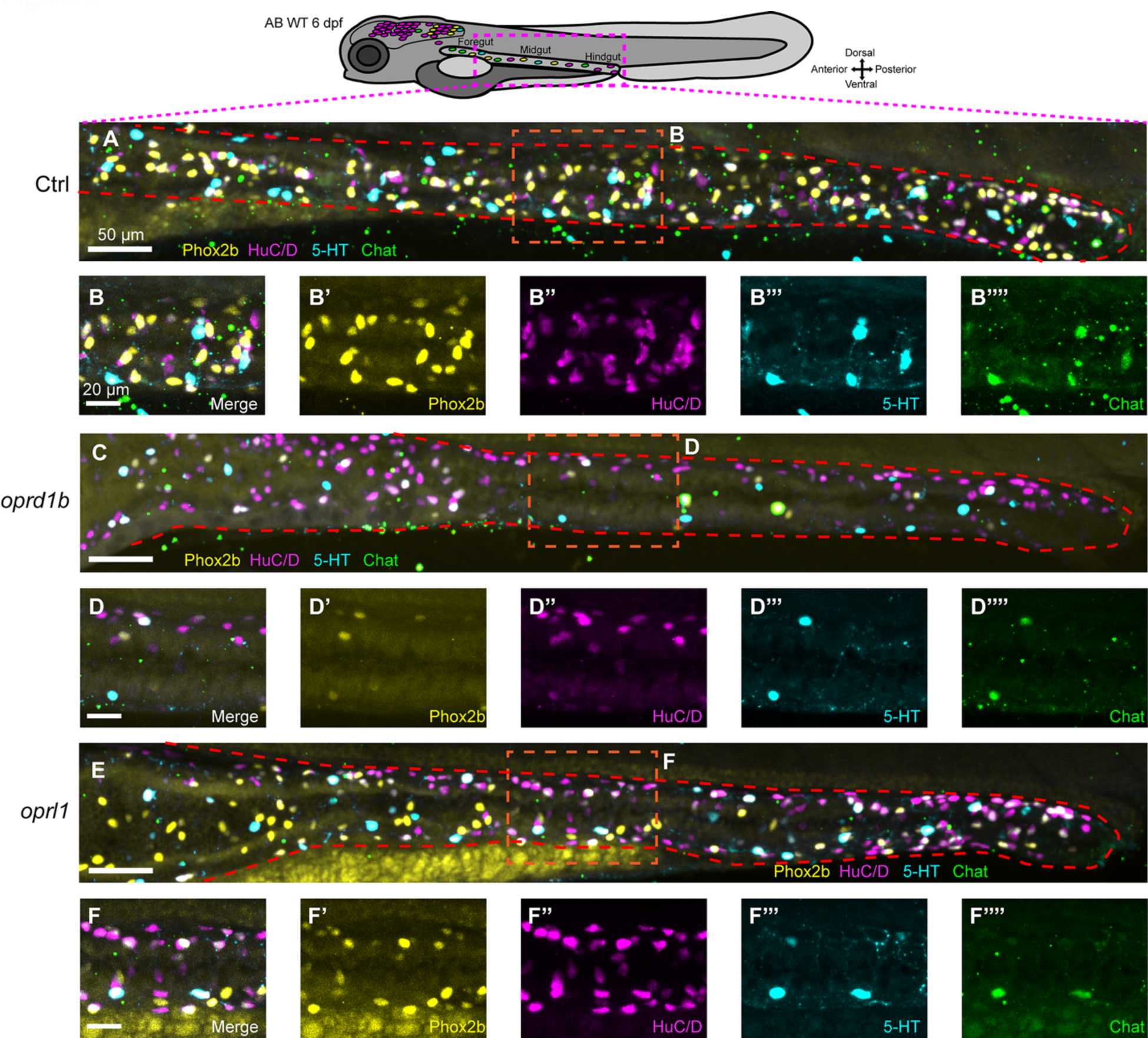
ENS neurochemical coding is altered in larval crispants for opioid receptor-encoding genes *oprd1b* and *oprl1*. (A, C, E) Confocal images show whole ENS after immunohistochemistry in 6 dpf control, and *oprd1b* and *oprl1* crispants, respectively. The targeted proteins were Phox2b (yellow), HuC/D (magenta), 5-HT (cyan) and Chat (green). Dashed red lines surround the gut. (B-B””, D-D””, F-F””) depict individual channels and magnification of the ENS midgut region to show the different marker proteins dissected by colors.

## Discussion

The robust synergy between scRNA-seq data and F0 CRISPR-based reverse genetics in the zebrafish model has allowed us to identify various novel genes involved in establishment of the ENS during development. This process involved bioinformatic identification of candidate genes from scRNA-seq data, drawing on differentially expressed genes and enrichment analysis. In addition, subsequent steps in the screen necessitating indel validations and culminating in high-throughput confocal imaging and analysis this proved to be notably swift, particularly when targeting an ENS colonization phenotype for identification. Overall, we focused on twelve candidate genes, two of which were positive controls already known to be required for zebrafish ENS development (*ret* and *gfra1a*). Among novel targets, we found that the genes *oprd1b*, *oprl1, rufy3, etv1, vipb, vgf, ache* and *flot1a,* when mutated caused significant ENS loss in crispants at 72 hpf (summarized in Table 1). Focusing our efforts on *oprd1b* and *oprl1,* we determined that inhibition of the opioid receptors encoded by these genes phenocopied their corresponding crispants, bringing to light the opioid pathway as a regulator of ENS formation.

Our screening approach bears resemblance to the study conducted by Gui et al. 2017, wherein they detected *de novo* mutations through exome sequencing of HSCR patients. Similar to our methodology, they utilized the Tg(-*8.3phox2bb*:Kaede) transgenic zebrafish (Harrison et al., 2014) and performed a comparative analysis between morpholino-mediated knockdown and CRISPR knockouts of six genes. Another recent ENS F0 screen was able to identify the role of ten transcription factors, finding alterations in the number of ENs, and in gut motility using Tg(*phox2bb*:GFP) embryos (Davidson et al., 2021). One of the features of our screen was the capability of identifying most of our candidate genes with alterations in the ENS. This accomplishment was likely due to streamlined analysis of the scRNA-seq sub-clustering and differential expression in combination with Metascape and STRING analyses, demonstrating the ability to rival and produce equivalent outcomes compared to the more intricate approaches (Parvez et al., 2021), for example where they targeted 188 genes to identify 16 genes that are important for the zebrafish embryonic heart.

Of the twelve candidate genes we screened, phenotyped (**Figure 2**, **Table 1**) and validated (**Figure 3**), two of them served as positive controls, having known knockout/knockdown zebrafish models. For *ret^wmr1^* and *ret^hu2486^* mutants, they present with HSCR-like phenotypes in larval fish (Baker et al., 2022; Heanue et al., 2016), and for *gfra1α* morphants, they had a reduction in the number of ENs, displaying hypoganglionosis (Shepherd et al., 2004). To our knowledge, for the additional ten genes there are no mutational models or phenotypes described in the ENS, highlighting the importance of our screen for illuminating these novel genes with different functions.

Overall, most of our targeted genes have at least one known function related to neurons (**Supplementary Figure 1** and **Table 1**) and had hypoganglionosis alterations in our screen (**Figure 2D**). Only *pbx3b* (Pre-B-cell leukemia transcription factor 3b) and *nos1* (Nitric oxide synthase) didn’t show clear alteration in ENS colonization. For the case of *pbx3b*, this could possibly be explained by a compensation effect, where *pbx3a* may compensate for loss of *pbx3b*. While not highly enriched, *pbx3a* is expressed within enteric neuronal populations of our scRNA-seq (Howard et al., 2021; data not shown). As well, *pbx3b* and *nos1* are expressed almost exclusively in sub-cluster 3 (**Figure 1B)** suggesting important functions for these two genes when neurons are maturing and/or differentiated (Howard et al., 2021). Looking to the future, it will be important to functionally validate each of our candidate screen hits, to unravel their specific roles during ENS development, as well as how they may fit within an enteric gene regulatory network.

Of note, the *oprd1b* and *oprl1* crispants, and opioid receptor inhibition, was sufficient to cause hypoganglionosis in our study, bringing to light the opioid pathway during ENS development. Moreover, when we assayed for changes in the neurochemical coding of ENs in 6 day old larvae, we found a drastic change in neuronal cell population composition suggesting an important role of opioid receptors during ENS development and neurogenesis. The opioid receptors have been studied greatly in adult ENS, with respect to the effect of opiates, synthetic opiates and endogenous opioid release, to understanding GI functions, such as motility and secretion (Galligan and Sternini, 2017; Wood and Galligan, 2004). In the mouse brain during development, it has been noted that exposure to morphine and the μ-opioid peptide receptor (MOR) agonist inhibits neural stem and progenitor cells (NSPCs) proliferation by slowing the cell cycle G2/M phase (Sargeant et al., 2008). It is possible a similar process in the developing ENS exists, where ENCCs may slow their proliferation along the gut following downregulation of their opioid receptors (figure 2D), or by inhibition of them (Figure 4B). Furthermore, in NSPCs it has been reported that different opioid receptors play a role in neural differentiation and that endogenous opioid systems modulate neural growth and development (Kibaly et al., 2019). Thus, the neurochemical phenotypes we observed in the opioid crispants, where the predominant cell population observed was HuC/D^+^ (Figure 5), may be due to a general compensatory increase in neural differentiation, or it may signify an alteration in EN subtype differentiation and/or distribution along the gut.

Ultimately, our zebrafish F0 CRISPR screen of the ENS was able to identify novel genes that are important during ENS development. This screen proved to be efficient by using a combination of scRNA-seq analysis, reverse CRISPR genetics and high content imaging. In the future, we can envision additional, in-depth functional analysis of each candidate gene, thereby increasing our understanding of ENS development in vertebrates, in normal and diseased states.

## Materials and Methods

### Zebrafish husbandry, and embryo larvae collection

This work was conducted in accordance with the Institutional Animal Care and Use Committee (IACUC) of Rice University. Embryos and larvae for all experiments were collected from controlled breeding of adult zebrafish for synchronous staging. All embryos were maintained at 28°C in standard E3 embryo medium until 24 h post fertilization (hpf), then were transferred to 0.003% 1-phenyl 2-thiourea (PTU)/E3 solution (Karlsson et al., 2001). Transgenic embryos used for this work include Tg(*−8.3phox2bb*:Kaede) (Harrison et al., 2014). Embryos and larvae were collected out of their chorions at the stage noted in each experiment.

### Transcriptomics and Enrichment analyses

Expression analysis was done using a zebrafish *sox10*:GFP single-cell RNA-seq atlas, Gene Expression Omnibus (GEO) database accession number GSE152906, and available on UCSC Cell Browser https://cells.ucsc.edu/?ds=zebrafish-neural-crest-atlas (Howard et al., 2021). Seurat was used to subset and generate sub-clusters from the 68-70 hpf neural neuronal groups 5 and 12. Data was visualized in feature plots, dot plots or violin plots. Differentially expressed gene markers from sub-cluster 3 (Supplementary Data1) were acquired by using the FindAllMarkers function (Howard et al., 2021). Metascape custom enrichment analysis (Zhou et al., 2019) of the sub-cluster 3 differentially expressed gene markers was done by selecting the top 800 genes using gene prioritization by evidence counting (GPEC) at 0.05 P-value cutoff using the Reactome gene sets from zebrafish. Networks for sub-cluster 3 were generated using STRING and Cytoscape (Shannon et al., 2003; Szklarczyk et al., 2023).

### CRISPR-Cas9 guide RNA design and Synthesis

Twelve sgRNAs were designed using the CRISPR design tool from Synthego (https://design.synthego.com/) using *Danio rerio* (GRCz11) genome and selecting top rank sgRNAs with high activity and minimal off targets (Doench et al., 2016) (Supplementary Table 2). Negative Control, Scrambled sgRNA #1, GCACUACCAGAGCUAACUCA (Synthego) was used as a negative control in pilot experiments.

### CRISPR-Cas9 microinjections and genotyping

Pools of 30 to 50 embryos fertilized from in-crossing Tg(−8.3*phox2bb*:Kaede) adults were injected at the one-cell stage in the yolk with a solution containing 100 picogram (pg) of gene specific sgRNA, 2 µM Cas9 NLS nuclease (Synthego) and Phenol red. A total of eight injected F0 larvae were dissociated, used in T7 endonuclease I activity assays (NEB E3321) as previously described (Baker et al., 2022), and the percentage of indels was determined. PCR pair of primers per gene used for theT7E1 activity (Supplementary table 1) targeted the specific sgRNA region for each gene and amplified regions between 200 to 300 bp. Experimental replicates were done at least 3 times per gene for the injections and the genotyping.

### *in vivo* high content semi-automated confocal microscopy

For the screen, 4 to 8 Tg(*−8.3phox2bb*:Kaede) 72 hpf F0 larvae were selected from each gene specific T7EI confirmed pool, and placed upon a 1% agarose cast made inside of a µ-Dish 35 mm, glass bottom dish (ibidi, 81158). This cast with space for 42 embryos was created from a 3D-printed stamp using a Formlabs Form1+ SLA printer (Kleinhans and Lecaudey, 2019). The embryos were anesthetized using 0.4% Tricaine and covered with a solution of 0.5% low melt temperature agarose dissolved in E3 media. Embedded fish were then covered in 1× PTU/E3 media supplemented with 0.4% Tricaine. Afterwards confocal imaging was performed in an Olympus FV3000 confocal and FluoView software (2.4.1.198), using an Olympus 10.0X objective (UPLXAPO10X) at a constant temperature of 28°C, maintained with an OKOLAB Uno-controller imaging incubator. The embedded fish were scanned in an automated fashion using the multi-area time-lapse software module (MATL). Z-stack images of the ENS were combined using the Fiji Image-J stitch plugin version 1.2 and then processed and exported in IMARIS image analysis software (Bitplane). Figures were prepared in Adobe Illustrator software.

### Hybridization chain reaction and whole mount immunohistochemistry

Hybridization chain reaction (HCR) and Whole Mount Immunofluorescence were done in accordance with previous described methods (Ibarra-García-Padilla et al., 2021; Uribe and Bronner, 2015). The HCR probes transcripts were synthesized by Molecular Instruments for *ret,* NM_181662.2; *gfra1a,* NM_131730.1; *oprl1*, NM_205589.2; *oprd1b, NM_131258.4*; *etv1*, XM_005157634.4; *elavl3,* NM_131449. The following primary antibodies were used: goat polyclonal IgG anti-Choline Acetyltransferase (ChaT, Millipore Sigma, AB144P, 1:500), rabbit polyclonal IgG anti-5-HT (serotonin, Immunostar, 20080, 1:250), mouse monoclonal IgG2b anti-HuC/D (Invitrogen Thermo Fisher, A-21271, 1:250), Mouse monoclonal IgG1 anti-Phox2b (B-11, Santa Cruz Biotechnology, SC-376997, 1:250). The following secondary antibodies were used from Invitrogen: Alexa Fluor 488 donkey anti-goat IgG (A-11055, 1:600), Alexa Fluor 405 goat anti-rabbit IgG (A-48254, 1:600), Alexa Fluor 647 goat anti-mouse IgG2b (A-21242, 1:600), Alexa Fluor 594 goat anti-mouse IgG1 (A-21125, 1:600). High content semi-automated confocal imaging and processing was done as described above.

### Zebrafish treatment with chemical inhibitors

LY2940094 (MedChemExpress, HY-114452), curcumin (sigma-aldrich, C1386) and DADLE (abcam, ab120673) master stocks were diluted in DMSO and then further diluted in 1xPTU/E3 medium to the required working concentration (10 µM). Eight Tg(−*8.3phox2bb*:Kaede) embryos per well were setup in a 24-well plate (Corning, CLS3527) with 1 ml of 1xPTU/E3. Drug was added at 48 hpf and incubated for 48 hpe (hours post exposure, until 96 hpf). Titration experiments were done to determine the working concentrations (data not shown). 48 hpf was chosen, as this is when ENCCs are actively migrating along the gut to colonize, as well as undergoing neurogenesis (Olden et al., 2008; Taylor et al., 2016). Larvae were extensively washed to remove treatments, then prepared and imaged by high content semi-automated confocal microscopy as described above. Experimental replicates were done 3 times with over 4 biological replicates each.

### Statistics

Statistical analysis was performed in GraphPad Prism (version 10.1.1). For comparisons, data was tested using two-tailed unpaired t-test and Ordinary one-way ANOVA, *P<0.05, n.s., non-significant (P<0.05).

## Acknowledgments

We want to express our sincere appreciation to Margarita Niño, Lucia J. Rivas and James J. Tallman, for their invaluable assistance in initiating experiments. Also, we want to thank Aubrey GA Howard IV, Phillip A Baker and Helen Folasade Adu for their insight, advice and technical assistance.

## Conflict of Interest

None.

## Funding Information

This study was supported by the National Institutes of Health grant R01DK124804 awarded to R.A.U., and by National Science Foundation grant 1942019 awarded to R.A.U. The funders had no role in study design, data collection and analysis, decision to publish, or preparation of the manuscript.

---

**Supplementary figure 1.**
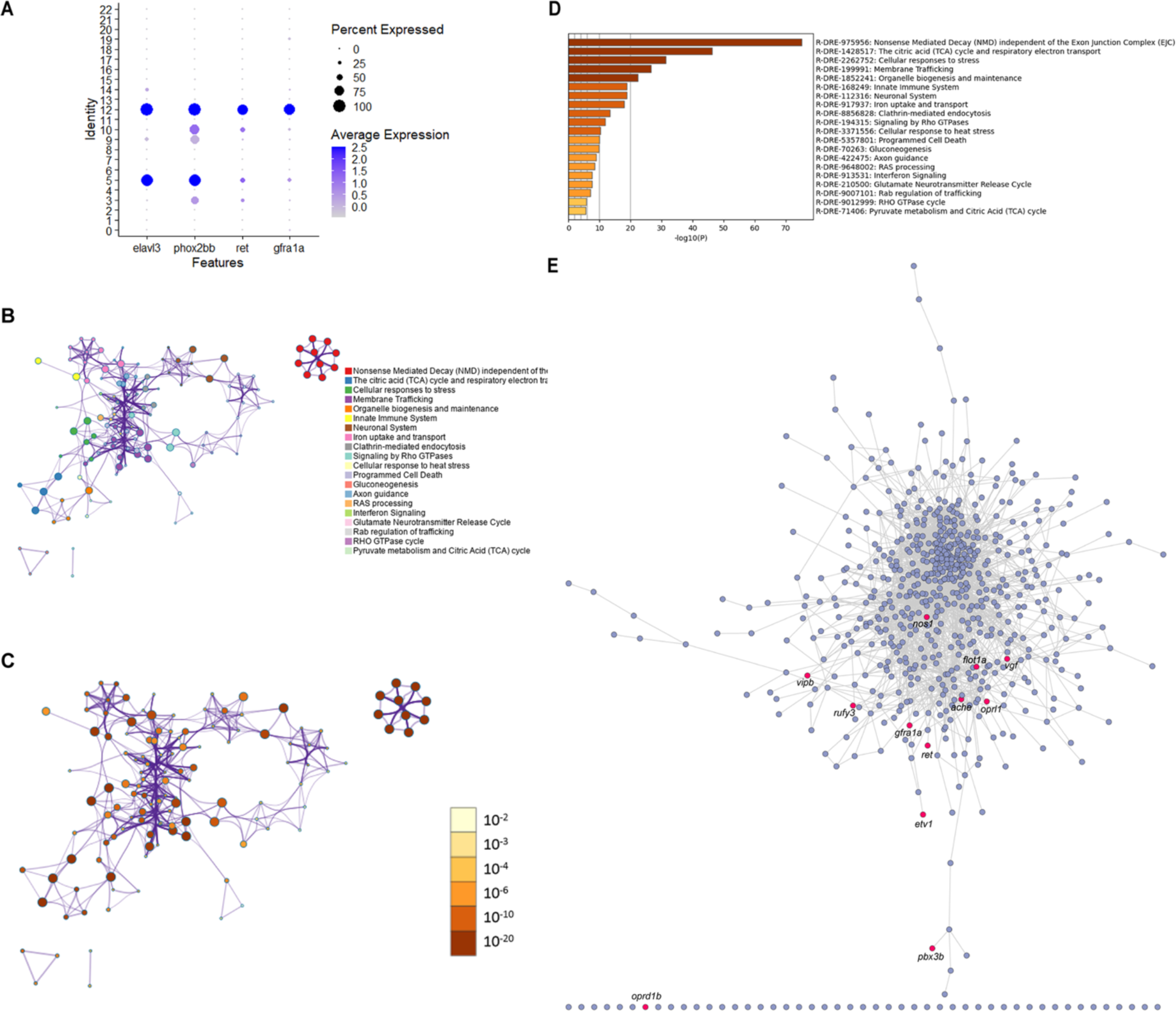
Selection of ENS genes for the CRISPR screen was based on scRNA seq analysis, Metascape functional enrichment and STRING analysis. (A) Dot plot depicts expression level of enteric neuron specific markers across individual clusters generated within the main 68-70 hpf tSNE *sox10*:GFP^+^ dataset (Howard et al., 2021). Clusters 5 and 12 prominently expressed enteric neuron markers. Dot size depicts the cell percentage for each marker and the color summarizes the average expression levels for each gene. (B, C) Metascape network of enriched terms from the reactome zebrafish gene set, colored by cluster (B) or by p-values (C), this network was based on the differentially expressed genes table from Seurat scRNA seq analysis of sub-cluster 3 (Supplementary Data 1). (D) Bar graph of top 20 enriched reactome zebrafish terms across input gene lists colored by p-values. (E) STRING network based on the complete sub-cluster 3 depicting in red color the selected genes for the CRISPR screen.

**Supplementary figure 2.**
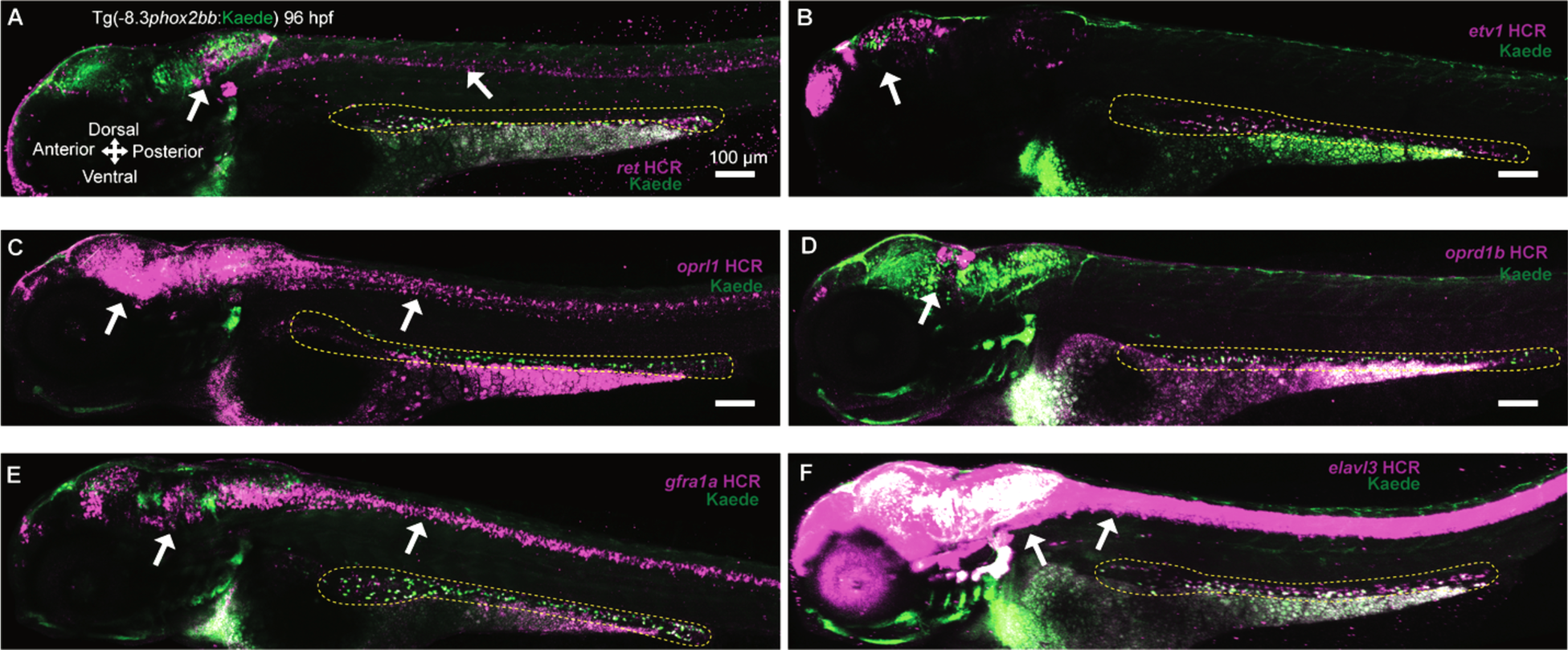
Whole body expression pattern of selected CRISPR screen ENS genes. Confocal images show HCR-assayed expression of *ret* (A), *etv1* (B), *oprl1* (C), *oprd1b* (F), *gfra1a* (E), and *elavl3* (F) in whole Tg(-8.3*phox2bb*:Kaede) larvae at the 96 hpf, revealing expression along the spinal cord and brain regions. Kaede signal is green and specific mRNA signal is magenta. White arrows depict the signals of the probes in the brain or spinal cord, dashed yellow lines surround the ENS.

**Supplementary Data 1. Differentially expressed genes of sub-cluster 3 from Seurat scRNA seq (Howard et al., 2021).**

**Supplementary Table 1. sgRNAs that target the different ENS candidate genes.**

## Notes

### Competing Interest Statement

The authors have declared no competing interest.

## References

Abe, H., Okazawa, M. and Nakanishi, S. (2011). The Etv1/Er81 transcription factor orchestrates activity-dependent gene regulation in the terminal maturation program of cerebellar granule cells. Proceedings of the National Academy of Sciences 108, 12497– 12502.

Baker, P. A., Ibarra-García-Padilla, R., Venkatesh, A., Singleton, E. W. and Uribe, Rosa. A. (2022). In toto imaging of early enteric nervous system development reveals that gut colonization is tied to proliferation downstream of Ret. Development 149, dev200668.

Bandla, A., Melancon, E., Taylor, C. R., Davidson, A. E., Eisen, J. S. and Ganz, J. (2022). A New Transgenic Tool to Study the Ret Signaling Pathway in the Enteric Nervous System. International Journal of Molecular Sciences 23, 15667.

Brosens, E., Burns, A. J., Brooks, A. S., Matera, I., Borrego, S., Ceccherini, I., Tam, P. K., García-Barceló, M.-M., Thapar, N., Benninga, M. A., et al. (2016). Genetics of enteric neuropathies. Developmental Biology 417, 198–208.

Char, R. and Pierre, P. (2020). The RUFYs, a Family of Effector Proteins Involved in Intracellular Trafficking and Cytoskeleton Dynamics. Frontiers in Cell and Developmental Biology 8,.

Choi, T.-Y., Choi, T.-I., Lee, Y.-R., Choe, S.-K. and Kim, C.-H. (2021). Zebrafish as an animal model for biomedical research. Exp Mol Med 53, 310–317.

Davidson, A. E., Straquadine, N. R., Cook, S. A., Liu, C. G. and Ganz, J. (2021). A rapid F0 CRISPR screen in zebrafish to identify regulators of neuronal development in the enteric nervous system. 2021.07.17.452230.

DiCello, J. J., Carbone, S. E., Saito, A., Pham, V., Szymaszkiewicz, A., Gondin, A. B., Alvi, S., Marique, K., Shenoy, P., Veldhuis, N. A., et al. (2022). Positive allosteric modulation of endogenous delta opioid receptor signaling in the enteric nervous system is a potential treatment for gastrointestinal motility disorders. American Journal of Physiology-Gastrointestinal and Liver Physiology 322, G66–G78.

Doench, J. G., Fusi, N., Sullender, M., Hegde, M., Vaimberg, E. W., Donovan, K. F., Smith, I., Tothova, Z., Wilen, C., Orchard, R., et al. (2016). Optimized sgRNA design to maximize activity and minimize off-target effects of CRISPR-Cas9. Nat Biotechnol 34, 184–191.

Douarin, N. M. L. and Teillet, M.-A. (1973). The migration of neural crest cells to the wall of the digestive tract in avian embryo. Development 30, 31–48.

Epstein, M. L., Mikawa, T., Brown, A. M. C. and McFarlin, D. R. (1994). Mapping the origin of the avian enteric nervous system with a retroviral marker. Developmental Dynamics 201, 236–244.

Fleming, M. A., Ehsan, L., Moore, S. R. and Levin, D. E. (2020). The Enteric Nervous System and Its Emerging Role as a Therapeutic Target. Gastroenterology Research and Practice 2020, e8024171.

Furness, J. B. (2008). The Enteric Nervous System. John Wiley & Sons.

Furness, J. B. and Stebbing, M. J. (2018). The first brain: Species comparisons and evolutionary implications for the enteric and central nervous systems. Neurogastroenterology & Motility 30, e13234.

Galligan, J. J. and Sternini, C. (2017). Insights into the Role of Opioid Receptors in the GI Tract: Experimental Evidence and Therapeutic Relevance. Handb Exp Pharmacol 239, 363–378.

Ganz, J. (2018). Gut feelings: Studying enteric nervous system development, function, and disease in the zebrafish model system. Developmental Dynamics 247, 268–278.

Geng, Z.-H., Zhu, Y., Li, Q.-L., Zhao, C. and Zhou, P.-H. (2022). Enteric Nervous System: The Bridge Between the Gut Microbiota and Neurological Disorders. Frontiers in Aging Neuroscience 14,.

Gui, H., Schriemer, D., Cheng, W. W., Chauhan, R. K., Antiňolo, G., Berrios, C., Bleda, M., Brooks, A. S., Brouwer, R. W. W., Burns, A. J., et al. (2017). Whole exome sequencing coupled with unbiased functional analysis reveals new Hirschsprung disease genes. Genome Biology 18, 48.

Harrison, C., Wabbersen, T. and Shepherd, I. T. (2014). In vivo visualization of the development of the enteric nervous system using a Tg(−8.3bphox2b:Kaede) transgenic zebrafish. genesis 52, 985–990.

Heanue, T. A., Boesmans, W., Bell, D. M., Kawakami, K., Berghe, P. V. and Pachnis, V. (2016). A Novel Zebrafish ret Heterozygous Model of Hirschsprung Disease Identifies a Functional Role for mapk10 as a Modifier of Enteric Nervous System Phenotype Severity. PLOS Genetics 12, e1006439.

Howard, A. G., IV, Baker, P. A., Ibarra-García-Padilla, R., Moore, J. A., Rivas, L. J., Tallman, J. J., Singleton, E. W., Westheimer, J. L., Corteguera, J. A. and Uribe, R. A. (2021). An atlas of neural crest lineages along the posterior developing zebrafish at single-cell resolution. eLife 10, e60005.

Howard, A. G. A., Nguyen, A. C., Tworig, J., Ravisankar, P., Singleton, E. W., Li, C., Kotzur, G., Waxman, J. S. and Uribe, R. A. (2022). Elevated Hoxb5b Expands Vagal Neural Crest Pool and Blocks Enteric Neuronal Development in Zebrafish. Frontiers in Cell and Developmental Biology 9,.

Hwang, W. Y., Fu, Y., Reyon, D., Maeder, M. L., Tsai, S. Q., Sander, J. D., Peterson, R. T., Yeh, J.-R. J. and Joung, J. K. (2013). Efficient genome editing in zebrafish using a CRISPR-Cas system. Nat Biotechnol 31, 227–229.

Ibarra-García-Padilla, R., Howard, A. G. A., Singleton, E. W. and Uribe, R. A. (2021). A protocol for whole-mount immuno-coupled hybridization chain reaction (WICHCR) in zebrafish embryos and larvae. STAR Protocols 2, 100709.

Karlsson, J., von Hofsten, J. and Olsson, P. E. (2001). Generating transparent zebrafish: a refined method to improve detection of gene expression during embryonic development. Mar Biotechnol (NY) 3, 522–527.

Kawai, K. and Takahashi, M. (2020). Intracellular RET signaling pathways activated by GDNF. Cell Tissue Res 382, 113–123.

Keck, S., Galati-Fournier, V., Kym, U., Moesch, M., Usemann, J., Müller, I., Subotic, U., Tharakan, S. J., Krebs, T., Stathopoulos, E., et al. (2021). Lack of Mucosal Cholinergic Innervation Is Associated With Increased Risk of Enterocolitis in Hirschsprung’s Disease. Cellular and Molecular Gastroenterology and Hepatology 12, 507–545.

Keren-Kaplan, T., Sarić, A., Ghosh, S., Williamson, C. D., Jia, R., Li, Y. and Bonifacino, J. S. (2022). RUFY3 and RUFY4 are ARL8 effectors that promote coupling of endolysosomes to dynein-dynactin. Nat Commun 13, 1506.

Khuansuwan, S., Barnhill, L. M., Cheng, S. and Bronstein, J. M. (2019). A novel transgenic zebrafish line allows for in vivo quantification of autophagic activity in neurons. Autophagy 15, 1322–1332.

Kibaly, C., Xu, C., Cahill, C. M., Evans, C. J. and Law, P.-Y. (2019). Non-nociceptive roles of opioids in the CNS: opioids’ effects on neurogenesis, learning, memory and affect. Nat Rev Neurosci 20, 5–18.

Kleinhans, D. S. and Lecaudey, V. (2019). Standardized mounting method of (zebrafish) embryos using a 3D-printed stamp for high-content, semi-automated confocal imaging. BMC Biotechnol 19, 68.

Kuil, L. E., Chauhan, R. K., Cheng, W. W., Hofstra, R. M. W. and Alves, M. M. (2021). Zebrafish: A Model Organism for Studying Enteric Nervous System Development and Disease. Frontiers in Cell and Developmental Biology 8,.

Kuil, L. E., Kakiailatu, N. J. M., Windster, J. D., Bindels, E., Zink, J. T. M., van der Zee, G., Hofstra, R. M. W., Shepherd, I. T., Melotte, V. and Alves, M. M. (2023). Unbiased characterization of the larval zebrafish enteric nervous system at a single cell transcriptomic level. iScience 26, 107070.

Lei, C., Sun, R., Xu, G., Tan, Y., Feng, W., McClain, C. J. and Deng, Z. (2022). Enteric VIP-producing neurons maintain gut microbiota homeostasis through regulating epithelium fucosylation. Cell Host & Microbe 30, 1417–1434.e8.

Martik, M. L. and Bronner, M. E. (2017). Regulatory Logic Underlying Diversification of the Neural Crest. Trends in Genetics 33, 715–727.

Montalva, L., Cheng, L. S., Kapur, R., Langer, J. C., Berrebi, D., Kyrklund, K., Pakarinen, M., de Blaauw, I., Bonnard, A. and Gosain, A. (2023). Hirschsprung disease. Nat Rev Dis Primers 9, 1–19.

Morarach, K., Mikhailova, A., Knoflach, V., Memic, F., Kumar, R., Li, W., Ernfors, P. and Marklund, U. (2021). Diversification of molecularly defined myenteric neuron classes revealed by single-cell RNA sequencing. Nat Neurosci 24, 34–46.

Mullick, M., Venkatesh, K. and Sen, D. (2017). d-Alanine 2, Leucine 5 Enkephaline (DADLE)-mediated DOR activation augments human hUCB-BFs viability subjected to oxidative stress via attenuation of the UPR. Stem Cell Research 22, 20–28.

Nagy, N. and Goldstein, A. M. (2017). Enteric nervous system development: A crest cell’s journey from neural tube to colon. Seminars in Cell & Developmental Biology 66, 94– 106.

Olden, T., Akhtar, T., Beckman, S. A. and Wallace, K. N. (2008). Differentiation of the zebrafish enteric nervous system and intestinal smooth muscle. genesis 46, 484–498.

Parvez, S., Herdman, C., Beerens, M., Chakraborti, K., Harmer, Z. P., Yeh, J.-R. J., MacRae, C. A., Yost, H. J. and Peterson, R. T. (2021). MIC-Drop: A platform for large-scale in vivo CRISPR screens. Science 373, 1146–1151.

Petrocelli, G., Pampanella, L., Abruzzo, P. M., Ventura, C., Canaider, S. and Facchin, F. (2022). Endogenous Opioids and Their Role in Stem Cell Biology and Tissue Rescue. International Journal of Molecular Sciences 23, 3819.

Rao, M. and Gershon, M. D. (2018). Enteric nervous system development: what could possibly go wrong? Nat Rev Neurosci 19, 552–565.

Sargeant, T. J., Day, D. J., Miller, J. H. and Steel, R. W. J. (2008). Acute in utero morphine exposure slows G2 / M phase transition in radial glial and basal progenitor cells in the dorsal telencephalon of the E15.5 embryonic mouse. European Journal of Neuroscience 28, 1060–1067.

Savidge, T. C. (2014). Importance of NO and its related compounds in enteric nervous system regulation of gut homeostasis and disease susceptibility. Current Opinion in Pharmacology 19, 54–60.

Seo, E.-J., Efferth, T. and Panossian, A. (2018). Curcumin downregulates expression of opioid-related nociceptin receptor gene (OPRL1) in isolated neuroglia cells. Phytomedicine 50, 285–299.

Shannon, P., Markiel, A., Ozier, O., Baliga, N. S., Wang, J. T., Ramage, D., Amin, N., Schwikowski, B. and Ideker, T. (2003). Cytoscape: A Software Environment for Integrated Models of Biomolecular Interaction Networks. Genome Res. 13, 2498–2504.

Sharkey, K. A. and Mawe, G. M. (2023). The enteric nervous system. Physiological Reviews 103, 1487–1564.

Shepherd, I. T., Pietsch, J., Elworthy, S., Kelsh, R. N. and Raible, D. W. (2004). Roles for GFRα1 receptors in zebrafish enteric nervous system development. Development 131, 241–249.

Statnick, M. A., Chen, Y., Ansonoff, M., Witkin, J. M., Rorick-Kehn, L., Suter, T. M., Song, M., Hu, C., Lafuente, C., Jiménez, A., et al. (2016). A Novel Nociceptin Receptor Antagonist LY2940094 Inhibits Excessive Feeding Behavior in Rodents: A Possible Mechanism for the Treatment of Binge Eating Disorder. J Pharmacol Exp Ther 356, 493– 502.

Szklarczyk, D., Kirsch, R., Koutrouli, M., Nastou, K., Mehryary, F., Hachilif, R., Gable, A. L., Fang, T., Doncheva, N. T., Pyysalo, S., et al. (2023). The STRING database in 2023: protein–protein association networks and functional enrichment analyses for any sequenced genome of interest. Nucleic Acids Research 51, D638–D646.

Talbot, J., Hahn, P., Kroehling, L., Nguyen, H., Li, D. and Littman, D. R. (2020). Feeding-dependent VIP neuron–ILC3 circuit regulates the intestinal barrier. Nature 579, 575–580.

Taylor, C. R., Montagne, W. A., Eisen, J. S. and Ganz, J. (2016). Molecular fingerprinting delineates progenitor populations in the developing zebrafish enteric nervous system. Developmental Dynamics 245, 1081–1096.

Uribe, R. A. and Bronner, M. E. (2015). Meis3 is required for neural crest invasion of the gut during zebrafish enteric nervous system development. Mol Biol Cell 26, 3728–3740.

Uyttebroek, L., Shepherd, I. T., Harrisson, F., Hubens, G., Blust, R., Timmermans, J.-P. and Van Nassauw, L. (2010). Neurochemical coding of enteric neurons in adult and embryonic zebrafish (Danio rerio). Journal of Comparative Neurology 518, 4419–4438.

Wang, Y., Qin, X., Han, Y. and Li, B. (2022). VGF: A prospective biomarker and therapeutic target for neuroendocrine and nervous system disorders. Biomedicine & Pharmacotherapy 151, 113099.

Wood, J. D. and Galligan, J. J. (2004). Function of opioids in the enteric nervous system. Neurogastroenterology & Motility 16, 17–28.

Zhan, Z., Ye, M. and Jin, X. (2023). The roles of FLOT1 in human diseases (Review). Molecular Medicine Reports 28, 1–18.

Zhou, Y., Zhou, B., Pache, L., Chang, M., Khodabakhshi, A. H., Tanaseichuk, O., Benner, C. and Chanda, S. K. (2019). Metascape provides a biologist-oriented resource for the analysis of systems-level datasets. Nat Commun 10, 1523.

